# Mice with Reduced PAR4 Reactivity show Decreased Venous Thrombosis and Platelet Procoagulant Activity

**DOI:** 10.1101/2024.10.14.617127

**Authors:** Elizabeth A. Knauss, Johana Guci, Norman Luc, Dante Disharoon, Grace H. Huang, Anirban Sen Gupta, Marvin T. Nieman

**Author notes:** Corresponding author: Marvin T. Nieman, Department of Pharmacology, Case Western Reserve University, 2109 Adelbert Road W309B, Cleveland, OH, 44106-4965, USA.

## Abstract

**Background:** Hypercoagulation and thrombin generation are major risk factors for venous thrombosis. Sustained thrombin signaling through PAR4 promotes platelet activation, phosphatidylserine exposure, and subsequent thrombin generation. A single-nucleotide polymorphism in PAR4 (rs2227376) changes proline to leucine extracellular loop 3 (P310L), which decreases PAR4 reactivity and is associated with a lower risk for venous thromboembolism (VTE) in a GWAS meta-analysis.

**Objective:** The goal of this study is to determine the mechanism for the association of rs2227376 with reduced risk for VTE in using mice with a homologous mutation (PAR4-P322L).

**Methods:** Venous thrombosis was examined using our recently generated PAR4-P322L mice in the inferior vena cava stasis and stenosis models. Coagulation and clot stability was measured using rotational thromboelastometry (ROTEM). Thrombin generating potential was measured in platelet-rich plasma. Phosphatidylserine surface expression and platelet-neutrophil aggregates were analyzed using flow cytometry.

**Results:** PAR4^P/L^ and PAR4^L/L^ had reduced incidence and size of venous clots at 48 hours. PAR4^P/L^ and PAR4^L/L^ platelets had progressively decreased phosphatidylserine in response to thrombin and convulxin, which led to decreased thrombin generation and decreased PAR4-mediated platelet-neutrophil aggregation.

**Conclusions:** The leucine allele in extracellular loop 3, PAR4-322L leads to fewer procoagulant platelets and decreased endogenous thrombin potential. This decreased ability to generate thrombin offers a mechanism for PAR4’s role in VTE highlighting a key role for PAR4 signaling.

## Introduction

Venous thrombosis is driven by three factors known as Virchow’s triad: blood flow stasis, endothelial activation, and hypercoagulation [1]. Clot development involves an interplay between immune cells, platelets, endothelial cells, and coagulation factors in a process known as immunothrombosis [2,3]. Neutrophils and platelets bind to activated endothelial cells and express a procoagulant surface that promotes coagulation and thrombin generation [4–6]. Thrombin is produced in massive amounts at the site of clot development via the coagulation cascade [7,8]. Its main functions are to convert fibrinogen to fibrin, an essential component of venous thrombi [9]. Thrombin also activates cells, predominantly endothelial cells and platelets, directly via protease-activated receptors (PARs) [10]. On human platelets, thrombin cleaves PAR1 and PAR4 [11]. PAR1 elicits a rapid transient signaling cascade, whereas PAR4 leads to prolonged signaling and is required for sustained platelet activation [12]. PAR4 requires more thrombin for activation than PAR1, and its unique signaling kinetics make it a promising therapeutic target to prevent excessive and pathological platelet activation. PAR4 cleavage by thrombin leads to platelet activation and aggregation [13]. PAR4 antagonists, like BMS-986120 and BMS-986141, show significant potential in animal models and early clinical trials with a low risk of bleeding and a high antithrombotic potential [14–16]. Given that PAR4-mediated platelet activation also promotes venous thrombosis in mice [17], it is worth further exploring PAR4 as a therapeutic target against platelet activation in thrombosis and inflammation.

Platelets are widely known for their essential roles in hemostasis and preventing excessive blood loss upon injury [18]. Platelets respond to vascular damage and inflammation by adhering to the activated endothelium and aggregating with each other [19]. After sustained activation they develop a procoagulant surface covered in phosphatidylserine that serves as a scaffold for the assembly of coagulation factors, leading to thrombin and fibrin generation to plug the broken vasculature [4]. Unchecked coagulation and excessive platelet activation leads to thrombosis. Platelets are known to promote arterial thrombosis and have been targeted for prevention [20]. Platelets also drive the early stages of venous thrombosis, but antiplatelets are not currently used to prevent venous events. Current preventative therapies for venous thrombosis and its complication pulmonary embolism, known collectively as venous thromboembolism (VTE), target coagulation to shut down thrombin generation and fibrin production. The downside of this approach is that normal hemostasis is also impaired, leading to bleeding complications. A targeted approach that reduces PAR4 activity would spare thrombin’s function on fibrinogen and other receptors like PAR1, allowing hemostasis to work unhindered.

Single nucleotide polymorphisms (SNPs) in PAR4 impact signaling and subsequent platelet function [21]. These SNPs impact patient response to antiplatelet therapies, and they can also be utilized as a tool in laboratories to further investigate the therapeutic potential of reducing PAR4 reactivity [22,23]. A SNP in PAR4 results in an amino acid switch at position 310 from a proline to a leucine (PAR4-P310L, rs2227376) [23]. The leucine at 310 is associated with reduced PAR4 signaling in cells, as measured by calcium signaling [23]. To determine how the PAR4-P310L variant impacts PAR4 function *in vivo*, we developed a mouse model with a homologous mutation at position 322 (PAR4-P322L). Mice with one (PAR4-322^P/L^, heterozygous) or two (PAR4-322^L/L^, homozygous) leucine alleles had a reduced platelet response to the PAR4-agonist peptide and thrombin. These mice also showed impaired arterial thrombosis and decreased clot stability [24].

Our lab, in collaboration with the INVENT consortium, also found that the leucine allele at PAR4-310 is associated with a 15% risk reduction in VTE [23]. Here, we used our mouse model to investigate the mechanism behind this association between PAR4 activity and venous thrombosis. PAR4-322^P/L^ and PAR4-322^L/L^ mice had significantly reduced venous thrombosis, and mice without PAR4 (PAR4-/-), were less likely to develop venous thrombi in the early stages. PAR4-P322L also resulted in decreased platelet procoagulant activity. We also showed that decreased PAR4 reactivity directly reduces platelet-neutrophil aggregation. These data demonstrate that PAR4 drives immunothrombosis by promoting platelet-neutrophil binding and inducing a procoagulant platelet phenotype that supports thrombin generation. Importantly, the PAR4-P322L mutation did not significantly impact hemostatic clot development as measured by rotational thromboelastometry, suggesting that PAR4 plays a more significant role in pathological platelet activation.

## Methods

### Animals

All animal experiments were performed in accordance with the approval from the Case Western Reserve University (CWRU) Animal Ethics Committee. The PAR4-P322L mutation was introduced into F2rl3 by inducing a C to T nucleotide switch using the CRISPR-Cas9 genome-editing system, as we have previously described [24]. PAR4 knockout mice (F2rl3-/-) were purchased from Mutant Mouse Regional Resource Centers. These mice are referred to as PAR4-322^P/P^ (wild type), PAR4-322^P/L^, PAR4-322^L/L^, or PAR4^-/-^. PAR4-P322L mice were bred via heterozygous crossings and genotyped via sanger sequencing.

### Venous Thrombosis Models

Venous thrombosis was modeled in male mice using the stasis and stenosis models of inferior vena cava (IVC) ligation [25,26]. For both models, mice were anesthetized using isoflurane (2%) and sterile technique was performed to open the abdomen and isolate the IVC. Visible back branches were cauterized, and the side branch was completely ligated when located below the renal vein. For the stasis model, the IVC was completely ligated with nylon suture (7-0, Ethilon). For the stenosis model, the suture was tied around a spacer (30g needle), which was removed immediately after ligation. For both surgeries, the peritoneum and outside skin were closed separately with nylon sutures, and 500-700 μL of saline was injected subcutaneously. Mice were subcutaneously given buprenorphine (0.05 mg/kg) immediately following surgery, then every 8-12 hours for pain management. Mice were placed on a heating pad daily and monitored for up to 48 hours.

### Clot Staining

Clots were harvested from mice at 48 hours and dissected from the surrounding tissue. Clots were added to tissue cassettes then fixed in 4% paraformaldehyde for 24 hours and stored in 70% ethanol at 4°C until processed. Fixed clots were embedded in paraffin and cut in 5 μM sections by the Tissue Resource Core at CWRU. H&E staining was also done by the tissue resource core at CWRU. After H&E staining, tissue blocks were sent to the imaging core at the Cleveland Clinic Lerner Research Institute for Carstairs staining.

### Evaluation of Blood Coagulation using ROTEM

Rotational thromboelastometry (ROTEM) was used to assess blood coagulation in whole blood from mice. 750 μL of whole blood was collected via IVC puncture into 10% sodium citrate (80 μL; 0.109 M). ROTEM was performed on a quad-channel computerized device (ROTEM Delta TEM International, Munich, Germany). Extrinsic pathway activation was assessed using the EXTEM assay. Within 2 hours of blood collection, 300 μL of each sample was placed in a disposable cuvette with an electric automated pipette connected to the ROTEM system. 20 μL of 0.2 M CaCl2 was added to recalcify the samples, followed by the addition of the EXTEM substrate (tissue factor) immediately before measurements began.The assay was run for 60 minutes. Clotting time (CT), clot formation time (CFT), maximum clot firmness (MCF), alpha angle, and the amplitudes at 10 (A10) and 20 (A20) minutes were determined [27,28]. CT is the time from the initiation of the reaction until the amplitude reaches 2 mm. CFT is the time between 2 and 20 mm of clot firmness. Alpha angle is the angle between the baseline and the clotting curve at 2 mm. MCF refers to the maximum amplitude of clot firmness. A10 and A20 represent the clot amplitude at 10 and 20 minutes, respectively.

### Thrombin Generation

Endogenous thrombin generating potential in platelet-rich plasma (PRP) was assessed using the TechnoThrombin® thrombin generation assay (TGA) kit (Diapharma, Cat #5006010). Whole blood was collected from mice via IVC puncture into sodium citrate. PRP from wild-type (PAR4-322P/P), PAR4-322^P/L^, PAR4-322^L/L^ or PAR4^-/-^ mice was prepared by centrifuging whole blood at 2300xg for 20 seconds. The plasma supernatant was carefully removed without disturbing the lower layer of red blood cells. Platelets were counted using a Coulter Counter (Beckman Coulter), and PRP was diluted to 2×10^8^ platelets/mL and added to a 96-well plate. TechnoThrombin® Reagent C High was used to initiate the reaction and a fluorogenic substrate was used to monitor thrombin generation at 37°C for 60 minutes. Fluorescence was measured immediately after addition of the reagent/substrate mixture with a Flexstation 3 (Molecular Devices) with SoftMax Pro software. The plate was read every minute for one hour at a 360 nm excitation and 460 nm emittance. A standard concentration of thrombin was diluted and measured over 10 minutes to create a standard curve for data analysis. Control samples containing high and low concentrations of human plasma were included to show daily variation in sample preparation and data collection.

### Procoagulant Platelet Assay

Phosphatidylserine (PS) exposure was detected on washed platelets using flow cytometry for FITC-tagged bovine lactadherin (Prolytix #BLAC-FITC). Whole blood was collected via IVC puncture into sodium citrate (0.109 M), diluted 1:4 in platelet wash buffer (10 mM sodium citrate, 150 mM NaCl, 1 mM EDTA, 1% dextrose) supplemented with PGI2, and centrifuged at 300xg for 5 minutes. The PRP supernatant was collected, diluted 4:1 in platelet wash buffer with PGI2, and centrifuged at 3000xg for 10 minutes. Pelleted platelets were resuspended in HEPES tyrode’s buffer (140 mM NaCl, 6.1 mM KCl, 2.4 mM MgSO_4_.7H2O, 1.7 mM Na_2_HPO_4_, 5.8 mM Na-HEPES), counted, then diluted to 1×10^8^ platelets/mL. 20 μL of diluted platelets were incubated with 5 μL convulxin, and 5 μL of either thrombin (Prolytix, BCT-1020) or PAR4-agonist peptide (AYPGKF). The final concentration of agonist in each reaction was 1 nM convulxin and 200 μM AYPGKF, 1 nM thrombin, or 5 nM thrombin. Following activation, platelets were labeled with 12 μg/mL of APC-tagged CD41a (eBioMWReg30, invitrogen #17-0411-80) and 5 μg/mL FITC-Lactadherin for 15 minutes in the dark. Samples were fixed with 2% paraformaldehyde and analyzed on a BD LSRII flow cytometer. Stimulation with a calcium ionophore (A23187, 15 μM) was included for each sample to define the positive population (data not shown).

### Platelet-Neutrophil Aggregates

Platelet-neutrophil aggregates were isolated as described previously [29]. Whole blood was collected via IVC puncture and diluted 1:10 in M199 supplemented with 100 U/mL heparin. White blood cells were labeled with PE-CD45 (clone 30-F11, invitrogen #12-0451-82), neutrophils with FITC-Ly6G (clone 1A8, invitrogen #11-9668-82), and platelets with APC-CD41a (eBioMWReg30, invitrogen #17-0411-80). Diluted whole blood was labeled and activated for 15 minutes at 37°C in the dark. BD FACs lysing solution was added for 15 minutes to lyse red blood cells and fix the remaining cells. Samples were centrifuged at 800xg for 5 minutes, and the pelleted cells were resuspended in phosphate-buffered saline. Platelet neutrophil-aggregates were analyzed on a BD LSRII flow cytometer at the Case Cytometry and Imaging Microscopy core.

## Results

### PAR4-P322L reduces venous thrombosis

The single nucleotide polymorphism at rs2227376 results in a C to T base substitution leading to an amino change from proline to leucine at position 310 in extracellular loop 3 of PAR4. Individuals with the minor allele, leucine, have a 15% risk reduction in VTE. To study the impact of this substitution, we generated a mouse model with the homologous mutation, PAR4-P322L [24]. These mutant mice display impaired platelet activation and aggregation, as well as delayed arterial thrombosis [24].

To determine whether PAR4-P322L impacts venous thrombosis, our initial experiments used the complete stasis model of inferior vena cava (IVC) ligation. PAR4-322^P/P^ (wild type) mice develop fully formed thrombi by 48 hours following complete IVC ligation. The IVC containing the thrombus was harvested and weighed (**Figure 1A**). The average weight of IVCs without thrombi (13.6 mg, data not shown) was subtracted to give the weight of the thrombus alone. Thrombi from PAR4-322^P/P^ mice averaged 17.7 ± 4.2 mg. Thrombi from PAR4-322^P/L^ mice were 8.4 ± 5.1 mg (p=0.004), while the sizes of clots from PAR4-322^L/L^ mice had a wide range that averaged 12.5 ± 8.5 mg. Thrombi from PAR4 knockout mice (PAR4^-/-^) were 8.1 ± 4.5 mg, significantly smaller than wild-types (p=0.002).

**Figure 1.**
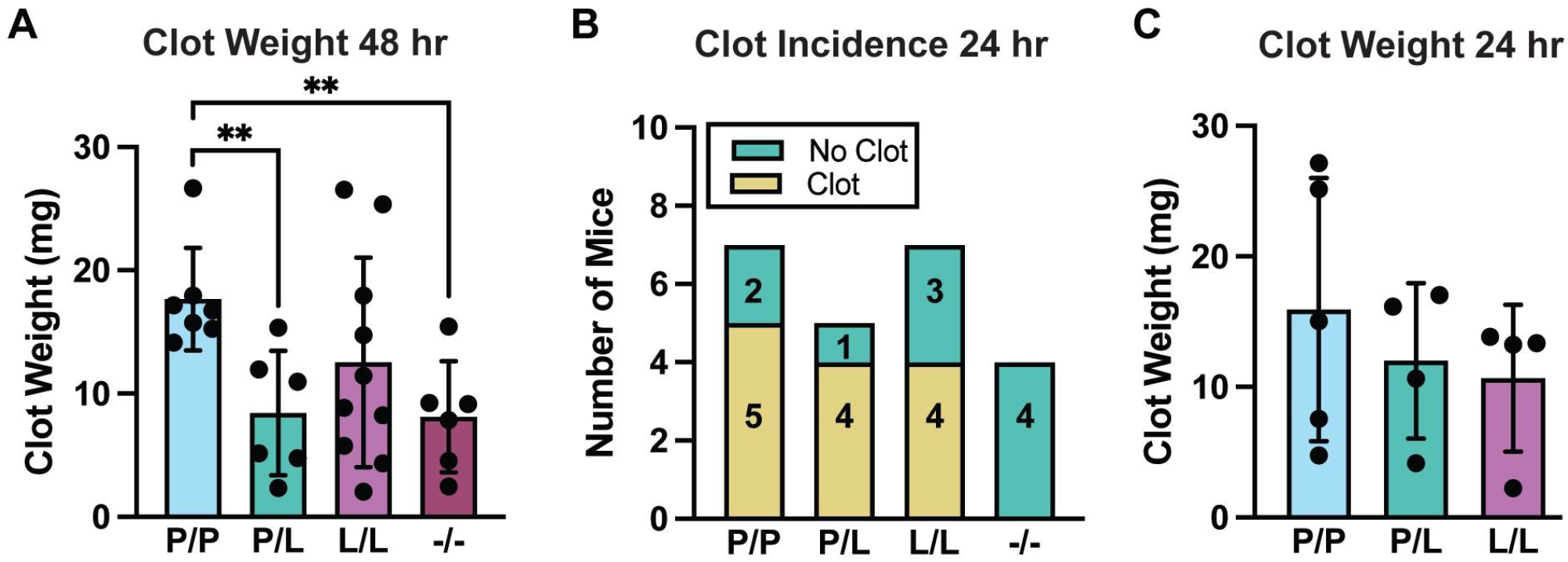
PAR4-P322L Reduces Venous Thrombosis after Complete IVC Stasis. **(A)** Venous clots within the inferior vena cava (IVC) were harvested from wild-type, heterozygous PAR4-322^P/L^, homozygous PAR4^L/L^, and global PAR4^-/-^ mice 48 hours after complete IVC ligation. **(B)** Clot incidence and **(C)** thrombus weight were also recorded 24 hours after IVC ligation. Dots represent individual mice.**p<0.01

Platelets have critical roles in the early stages of venous clot development [30]. To determine the impact of PAR4 at the initiation phase, we examined the time course of thrombus growth in PAR4-322^P/P^ mice at 12, 16, 20, and 24 hours. At 12, 16, and 20 hours, small clots formed at an incidence of ≤ 50% (data not shown). By 24 hours, PAR4-322^P/P^ mice developed both large and small thrombi more than half of the time (71% clot incidence) (**Figure 1B**). PAR4-322^P/L^ and 322^L/L^ mice similarly developed clots at 80% and 57% incidence, respectively. PAR4^-/-^ mice, however, were unable to develop clots by 24 hours. The thrombi that developed by 24 hours were 15.9 ± 10.1 mg for PAR4-322^P/P^, 12 ± 6 mg for PAR4-322^P/L^, and 10.7 ± 5.6 mg for PAR4-322^P/L^ (**Figure 1C**). Altogether, reducing or removing PAR4 activity decreased IVC thrombosis induced by complete venous stasis.

Partial stenosis of the IVC maintains residual blood flow and triggers inflammation at the site of clot development [26,31]. A major difference is the variability in resulting thrombus size and incidence compared to the stasis model. The reported variation is 65-100% thrombus occurrence at 48 hours. Wild-type mice developed clots 73% of the time (**Figure 2A**). 67% of PAR4-322^L/L^ mice and 40% of 322^P/L^ mice developed thrombi by 48 hours. The clots that formed in PAR4-322^P/P^ (WT) mice were 21.0 ± 6.2 mg (**Figure 2B**). Those from PAR4-322^P/L^ and PAR4-322^L/L^ mice were 8.8 ± 4.5 mg and 6.2 ± 4.8 mg (p=0.01), respectively. Overall, reducing PAR4 reactivity had an even greater impact on IVC stenosis than complete stasis.

**Figure 2.**
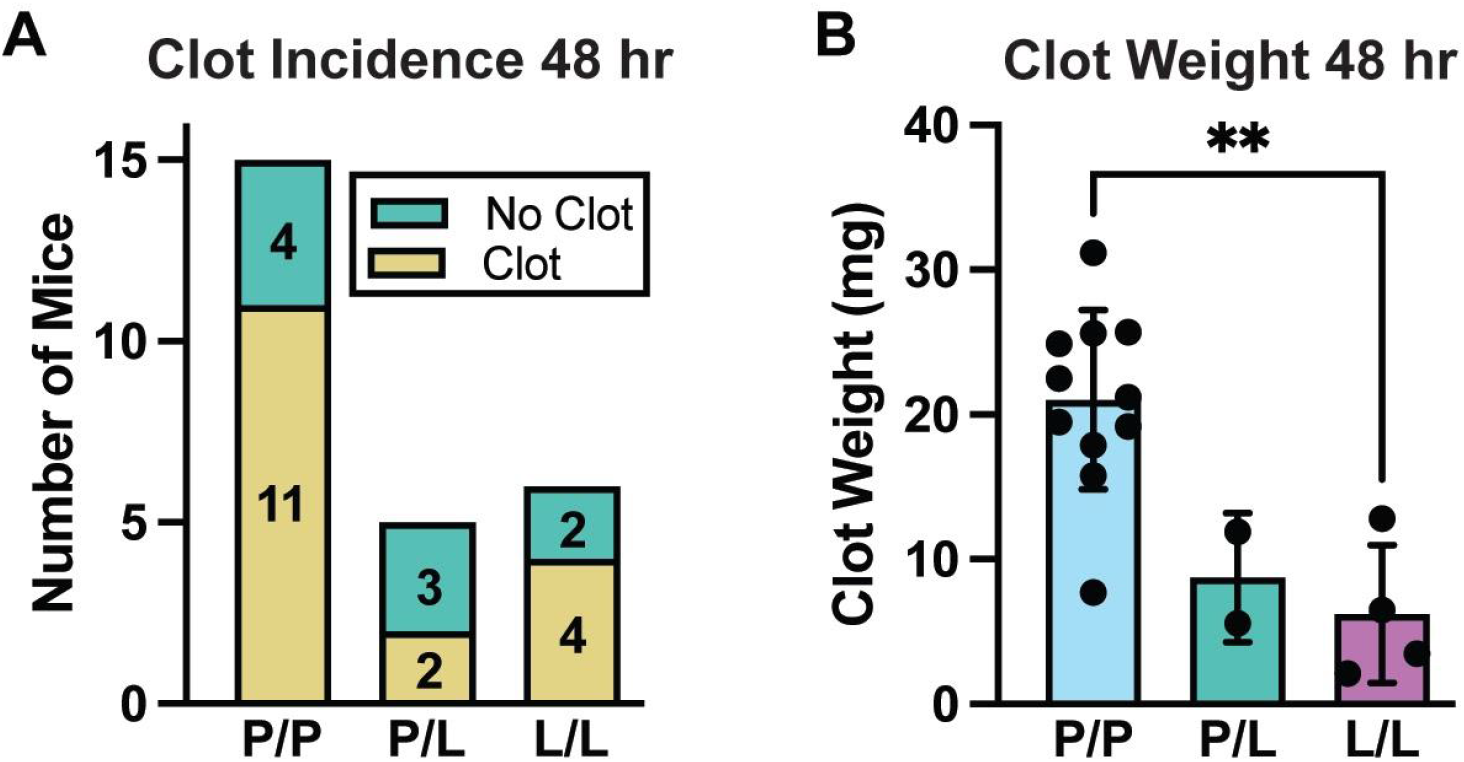
PAR4-P322L Reduces Venous Thrombosis after Partial IVC Stenosis. **(A)** The presence or absence of a thrombus was recorded 48 hours after IVC stenosis. **(B)** Clots were also excised from the surrounding tissue and weighed. Dots represent individual mice. *p<0.05

Clots that developed after IVC stenosis were processed for H&E and carstairs staining to observe any differences in clot composition or fibrin deposition, respectively. Both the PAR4-P322L mice and wild-types developed clots with layers of fibrin and white blood cells as well as lines of platelets [30,32].These characteristics are similar to those that define human thrombi, and indicate features that are essential to forming and stabilizing venous clots. H&E staining showed that while the clots from PAR4-322^P/L^ and PAR4-322^L/L^ mice were noticeably smaller than from wild-types, their composition was comparable (**Figure 3 panels A** and **B**). Thrombi from PAR4-P322L mice also showed a similar morphology to wild-types after carstairs staining (**Figure 3 panels C** and **D**). Thus, while the size of the clots decreases with PAR4 reactivity, the overall morphology and fibrin deposition remains the same.

**Figure 3.**
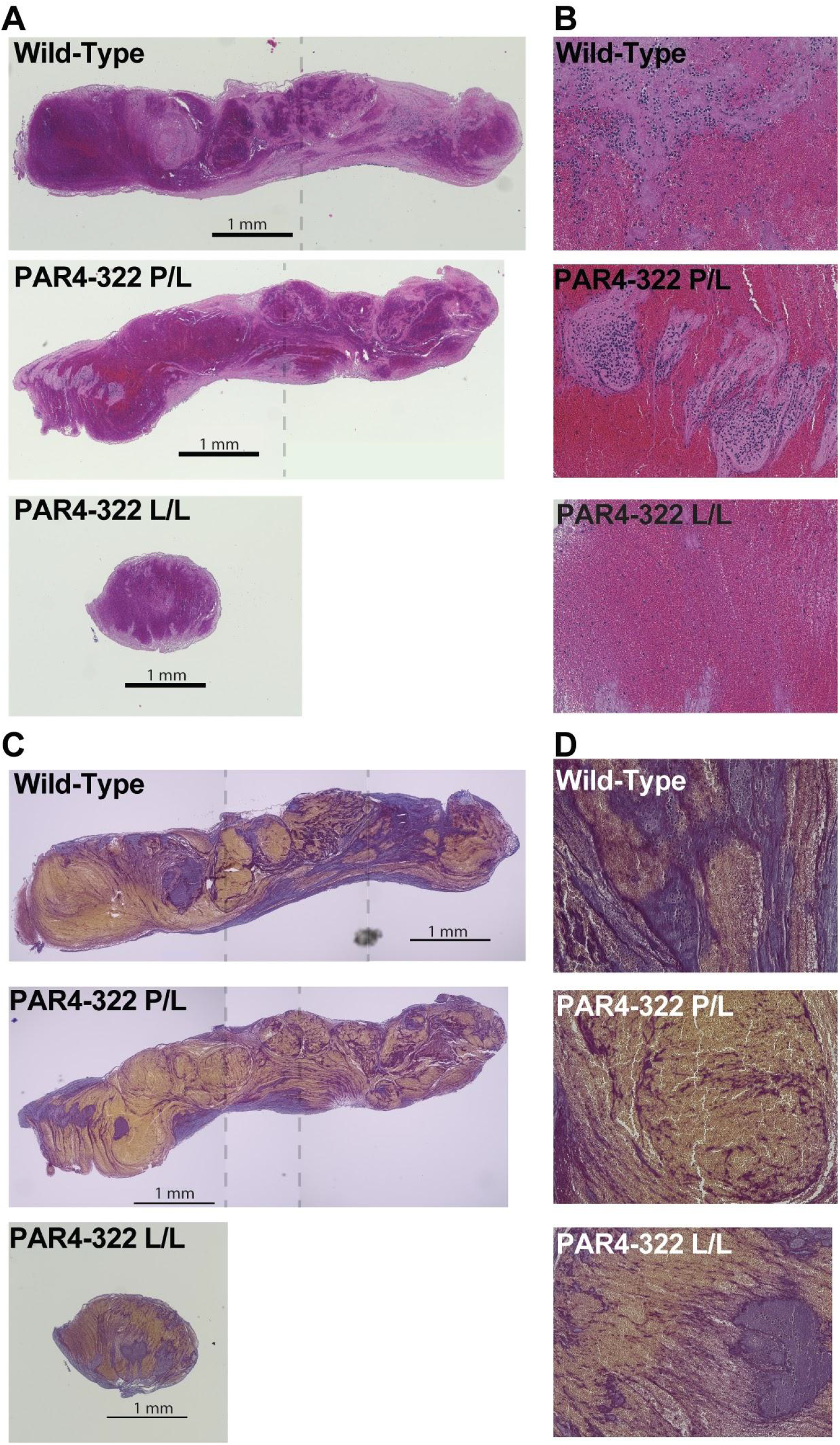
PAR4-P322L Mice have Smaller Thrombi after IVC Stenosis. Venous clots were excised from the surrounding tissue, embedded in paraffin, and sectioned for **(A)** haematoxylin and eosin (**A** and **B**) or carstairs (**C** and **D**) staining. Images were taken at 4x magnification and stitched together when necessary to display the entire clot (panels **A** and **C**). The stitching is indicated with the vertical dotted line. Higher resolution images (200x) are presented in panels **B** and **D**.

### Lack of PAR4 delays clot formation in ROTEM

We next used rotational thromboelastometry (ROTEM) to investigate the clot stability in whole blood from PAR4-322^P/P^, PAR4-322^P/L^, PAR4-322^L/L^, and PAR4^-/-^ mice. While ROTEM is traditionally used to measure hemostatic function, here we are specifically using it to measure tissue factor-induced coagulation using EXTEM. Thromboelastograms were developed for each sample over 60 minutes and clotting time (CT), clot formation time (CFT), maximum clot firmness (MCF) were determined from these traces (**Figure 4**). Clotting time (CT), also known as the coagulation time, indicates the initial activation of the coagulation cascade via tissue factor (**Figure 4B**) [27]. CT was unchanged in blood from mutant PAR4-P322L mice and global PAR4^-/-^ mice in comparison to wild-types, averaging around 40 seconds for each genotype. Clot formation time (CFT) was also unchanged in blood from both PAR4-322^P/L^ and 322^L/L^ mice (**Figure 4C**). CFT reflects the kinetics of clot formation and predominantly depends on platelet function, thrombin generation, and fibrin accumulation [27]. Blood from PAR4^-/-^ mice showed a pronounced increase in CFT and averaged 373 ± 93.9 seconds compared to wild-types at 41 ± 2.7 seconds (p < 0.0001). The MCF was unchanged across genotypes indicating that PAR4 does not impact the overall strength of the clot (**Figure 4D**) [27]. The rate of fibrin formation (alpha angle), and clot firmness (amplitude) at early time points were not different between PAR4-322^P/P^, PAR4-322^P/L^, PAR4-322^L/L^ mice, however all decreased in PAR4-/- mice (**Figure 4E-G**). Altogether, the allele at position 322 did not impact clot development in ROTEM stimulated with tissue factor. However the absence of PAR4 significantly delayed clot formation.

**Figure 4.**
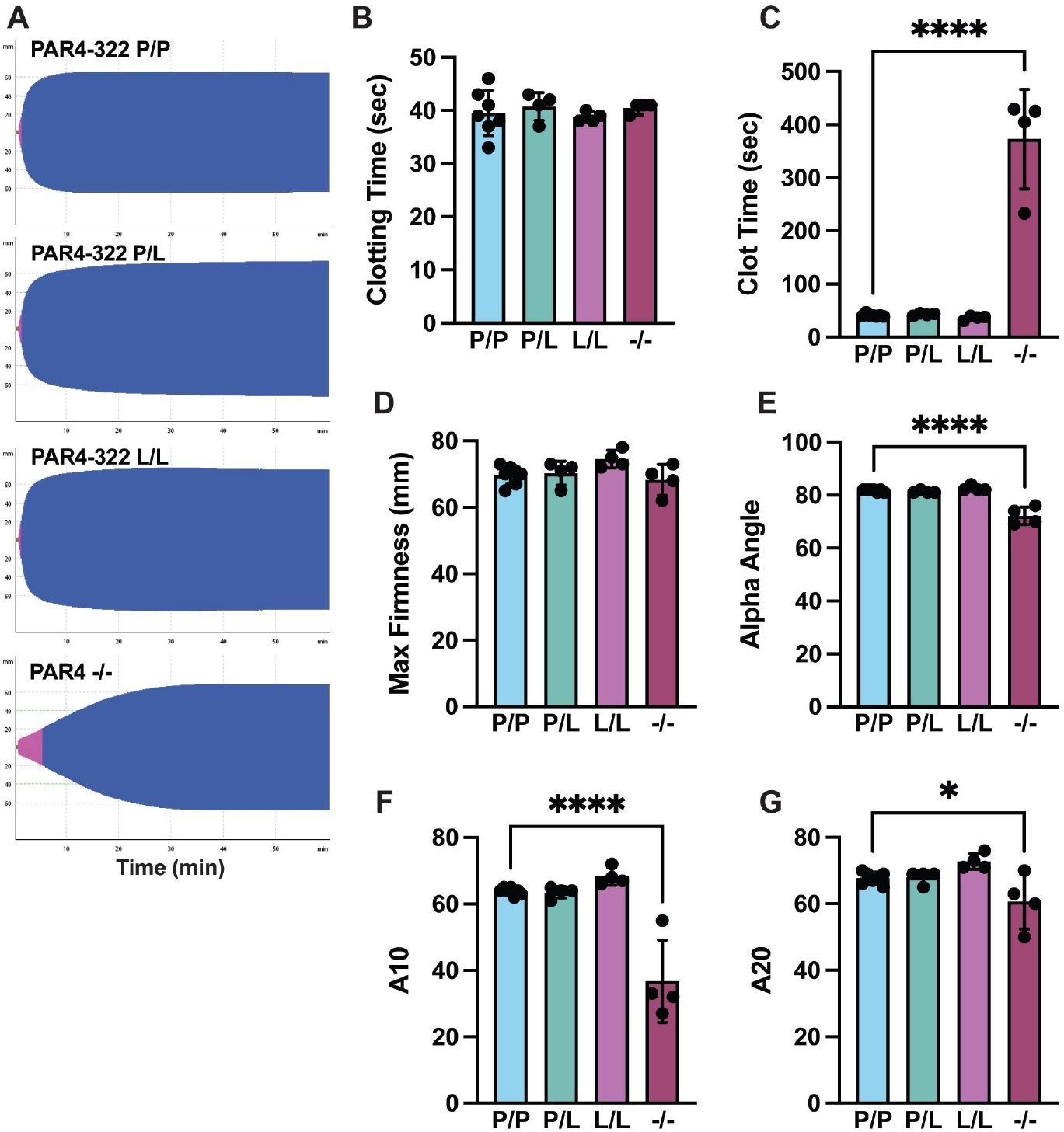
Blood from PAR4^-/-^ Mice Shows Impaired Clot Formation. ROTEM was performed to measure coagulation parameters by generating a thromboelastogram. **(A)** representative traces for each genotype are shown. **(B)** Clotting time, **(C)** clot formation time, **(D)** maximum clot firmness, **(E)** alpha angle, and the amplitudes at **(F)** 10 and **(G)** 20 minutes were extracted from the ROTEM traces. Dots represent individual mice. *p<0.05, ****p<0.0001.

### PAR4-P322L decreases phosphatidylserine exposure and thrombin generation

Thrombin is produced in massive amounts at the site of venous thrombosis where it activates platelets via PAR4 and cleaves fibrin to crosslink and stabilize venous clots [33]. At high concentrations of strong agonists like thrombin and collagen, platelets will adopt a procoagulant phenotype that serves as a platform for thrombin generation [5] [34,35]. We tested thrombin generating potential of platelet-rich plasma (PRP) from PAR4-322^P/P^, PAR4-322^P/L^, PAR4-322^L/L^ and PAR4^-/-^ mice for 60 minutes. (**Figure 5A**). The total thrombin generated (area under the curve), peak thrombin, and time to peak thrombin all showed a similar downward trend that correlated with PAR4 reactivity (**Figure 5B-D**). All parameters were significantly decreased in PRP from PAR4^-/-^ mice. Taken together, thrombin generating potential concomitantly decreased as PAR4 reactivity decreased.

**Figure 5.**
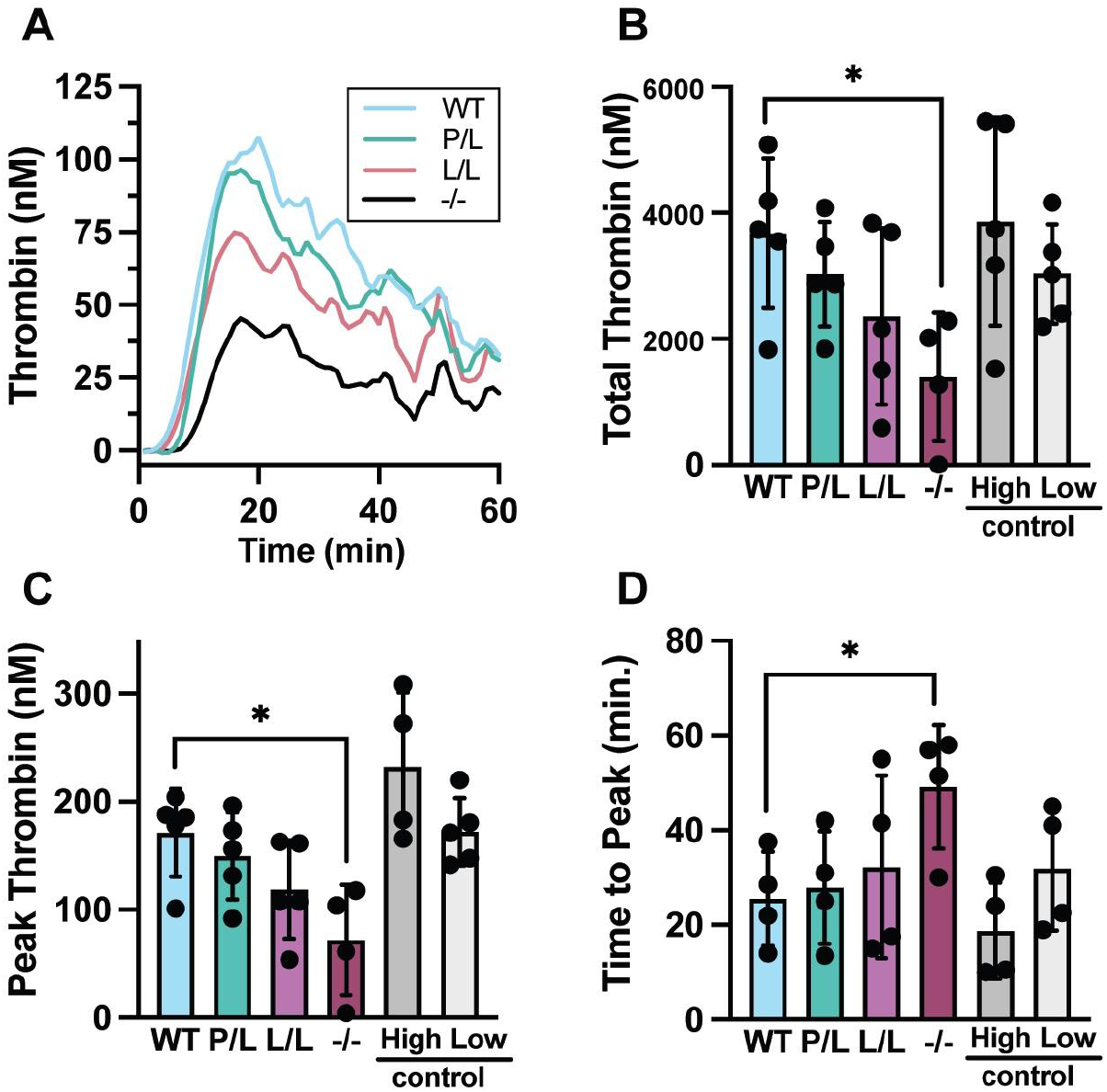
PAR4-P322L Decreases Platelet-Mediated Thrombin Generation. Platelet rich plasma (PRP) was obtained from wild-type, PAR4-322^P/L^, PAR4-322^L/L^, and PAR4^-/-^ mice and assessed for thrombin generating potential **(A-D). (A)** Thrombin generation curves were created using a fluorogenic thrombin substrate over 1 hour. **(B)** The area under the curve for each genotype was compared to the provided control human plasma (control high and control low) and interpreted as the total thrombin generated over that hour. **(C)** The peak concentration of thrombin generated and **(D)** the time it took to reach that peak was also recorded.

Platelet-mediated thrombin generation requires a procoagulant cell surface coated with phosphatidylserine (PS). We used FITC-labeled lactadherin to measure PS exposure on washed platelets. Platelets were stimulated with convulxin in combination with thrombin or the PAR4-agonist peptide (PAR4-AP), AYPGKF (**Figure 6)**. When stimulated with convulxin (1 nM) combined with AYPGKF (200 μM), 7.4% of PAR4-322^P/P^ platelets were PS-positive compared to 4.8% for PAR4-322^P/L^ platelets. Platelets from PAR4-322^L/L^ and of PAR4^-/-^ mice did not respond at this dose. Platelets from both PAR4-322^P/L^ and 322^L/L^ had less PS exposure on the surface following stimulation with convulxin and 1 or 5 nM thrombin. The PAR4^-/-^ platelets did not respond to thrombin as expected. These data indicate that the leucine allele at position 322 negatively impacts platelet PS exposure and subsequent thrombin generation, a phenotype that is amplified in platelets deficient in PAR4.

**Figure 6.**
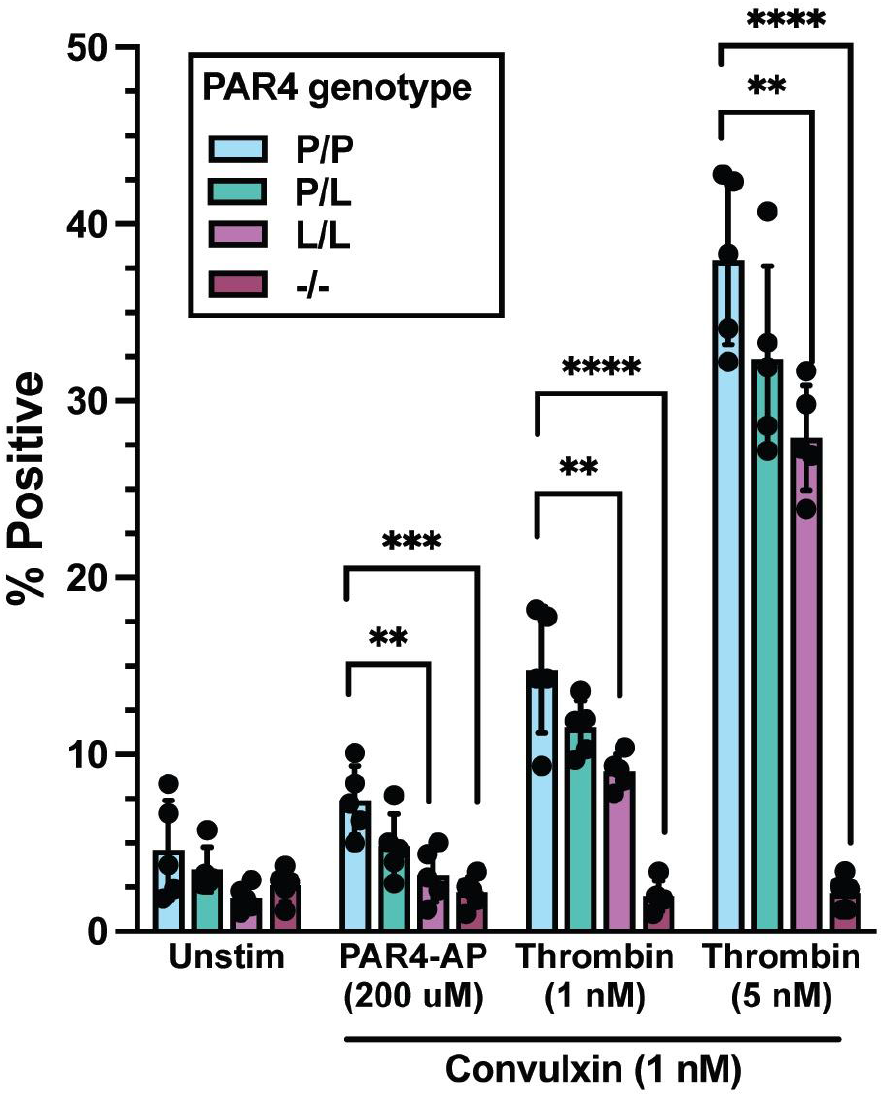
PAR4-P322L Reduces Platelet Phosphatidylserine Surface Expression. PS exposure was measured with flow cytometry using FITC-tagged Lactadherin. Washed mouse platelets were stimulated with convulxin (1 nM) in combination with 200 μM of the PAR4-AP (AYPGKF), 1 nM of thrombin, and 5 nM of thrombin. Lactadherin-positive events were recorded as the percentage of CD41-positive cells that were also positive for FITC. Dots represent individual mice. *p<0.05, **p<0.01, ***p<0.001, ****p<0.0001.

### PAR4-P322L decreases platelet-neutrophil complex formation

PAR4 activation is known to induce surface P-selectin exposure and platelet-neutrophil binding [36]. Additionally, procoagulant platelets preferentially activate neutrophils and promote thromboinflammation [5,37]. We have shown that mutant PAR4-P322L platelets have impaired P-selectin exposure in response to the PAR4-AP and thrombin [24]. Since platelet-neutrophil complexes (PNCs) are a risk factor for VTE, we examined the impact of PAR4-P322L on platelet-neutrophil aggregation. PNCs were defined as the percentage of CD45+, Ly6G+ cells that were positive for CD41. Complexes were measured at increasing concentrations of AYPGKF (50-400 μM) (**Figure 7A-D**). PAR4-322^P/P^ showed an increase in PNCs in response to 50 μM AYPGKF, while PAR4-P322L and PAR4^-/-^ did not respond (**Figure 7B**). PAR4-322^P/L^ mice responded to 200 μM, but the percentage of PNCs were significantly less than PAR4-322^P/P^ (**Figure 7C**). In contrast, PAR4^L/L^ and PAR4^-/-^ mice did not respond at the highest dose tested (400 μM) (**Figure 7D**). These data demonstrate that PAR4 reactivity is directly correlated to the formation of PNCs.

**Figure 7.**
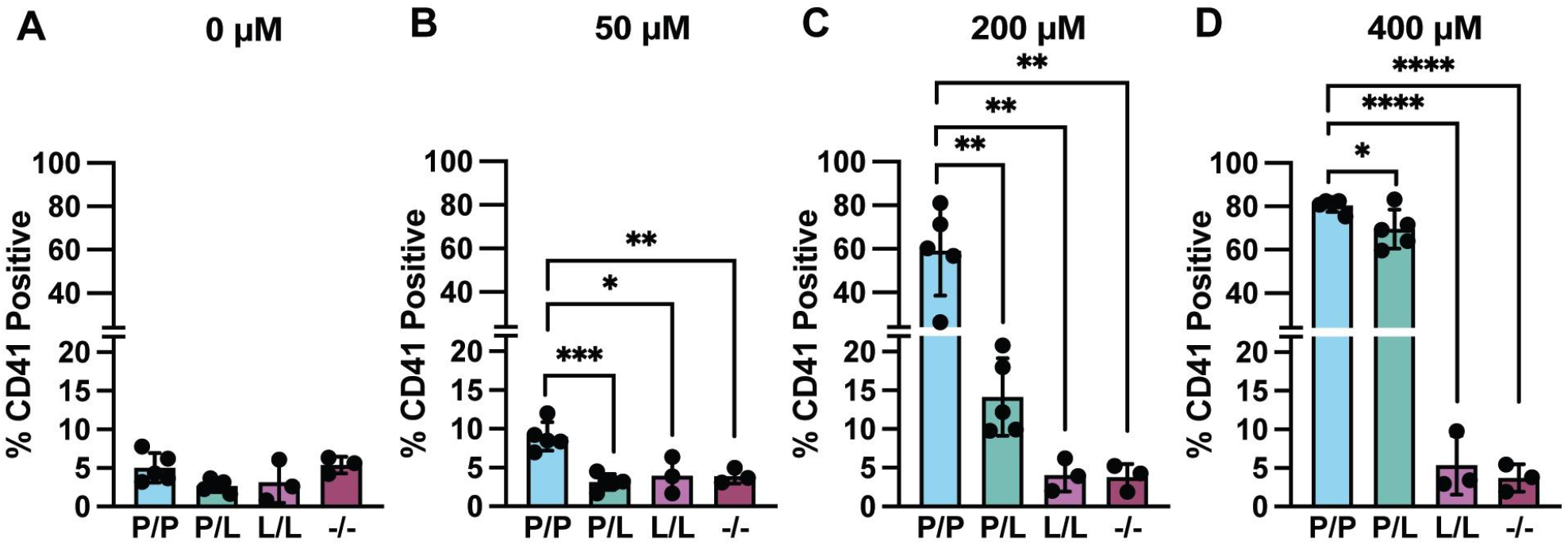
PAR4-P322L Decreases Platelet-Neutrophil Aggregation. Platelet-Neutrophil complexes were measured using flow cytometry to record the percentage of LyG+ neutrophils that also expressed CD41. Lysed whole blood from wild-type, PAR4-322^P/L^, PAR4-322^L/L^, and PAR4^-/-^ mice were incubated with with **(A)** no agonist, or **(B)** 50 μM, **(C)** 200 μM, or **(D)** 400 μM of the PAR4-agonist peptide (AYPGKF). *p<0.05, **p<0.01, ***p<0.001, ****p<0.0001.

## Discussion

The single-nucleotide polymorphism (SNP) that results in PAR4-Pro310Leu (rs2227376) lowers PAR4 activity and decreases PAR4-mediated platelet activation and arterial thrombosis [24]. The leucine allele at PAR4-310 is also associated with a 15% risk reduction in venous thromboembolism (VTE) [23]. Here, we used our mouse model of the PAR4-P310L variant (PAR4-P322L) to investigate the role of PAR4 in venous thrombosis. The PAR4-P322L mouse model has a point mutation at a site homologous to human PAR4-P310 and offers a way to study reduced PAR4 function *in vivo*. Here, we have shown that PAR4-P322L decreases platelet procoagulant activity and negatively impacts venous thrombosis.

We used the stasis model of inferior vena cava (IVC) thrombosis to mimic a completely occlusive thrombi in PAR4-322^P/P^ (wild-type), PAR4-322^P/L^, 322^L/L^, and PAR4^-/-^ mice. Complete IVC ligation is known for consistently inducing large thrombi with a low variability in size, making it a reliable technique for modeling acute thrombosis [31,38]. The PAR4-P322L mutation reduces PAR4 reactivity in response to thrombin [24]. These data suggest it also reduce platelet and endothelial cell activation in the early stages of thrombosis leading to smaller thrombi. By 48 hours, when the clots had fully formed, both PAR4-322^P/L^ and PAR4^-/-^ mice had significantly smaller venous thrombi than wild-types. Interestingly, PAR4-322^L/L^ mice developed thrombi with a much greater variability in size, with some as large as wild-types and others as small as PAR4^-/-^ (**Figure 1A**). When clots were harvested at 24 hours, PAR4^-/-^ were unable to develop venous thrombi, and PAR4-322^L/L^ mice only formed clots 50% of the time (**Figure 1B**). This suggests that PAR4 is driving the early stages of VTE when immune cells interact with coagulation factors to generate fibrin and initiate thrombosis [30].

Venous thrombi that develop from the stasis model of IVC ligation are similar in structure to those from humans, however they do not arise in the same manner. Complete IVC stasis predominantly produces clots through endothelial damage and tissue factor (TF) activation. The stenosis model of partial IVC ligation produces clots through a mechanism that is more dependent on immune cell activation [31].

The stenosis model avoids extreme endothelial damage, and mimics less occlusive thrombi that are more commonly seen in humans [26]. A downside of this model is that thrombosis is less likely to occur, but it allows a more sensitive method for analyzing thrombosis incidence. Indeed, we saw that both PAR4-322^P/L^ and PAR4-322^L/L^ mice developed fewer venous thrombi at 48 hours after partial IVC ligation (**Figure 2A**). Additionally, the clots that developed in the mutant mice were significantly smaller than wild-types (**Figure 2B**). The difference between the PAR4-P322L and wild-type mice was more pronounced in the stenosis model compared to complete stasis. The loss of PAR4 activity showed the greatest impact in the stasis model at early time points, suggesting a critical role in immunothrombosis and clot initiation. Unlike complete stasis, partial stenosis retains residual blood flow and triggers clot development through local inflammation and immunothrombosis [26]. Given that PAR4 is most highly expressed in this system on platelets and endothelial cells, it is likely that these cells are mediating PAR4’s role in thrombosis. Additionally, our data is consistent with others that showed platelet-specific knockout mice had decreased venous thrombus size and incidence after IVC stenosis [17]. Knowing that the PAR4-P322L mutation directly impacts platelet activation, we used our mouse model to further explore PAR4-mediated platelet activation in venous thrombosis.

Venous clots that form from complete ligation of the inferior vena cava are heavily driven by TF expressed upon endothelial damage [25]. TF is predominantly known for triggering extrinsic coagulation upon vascular injury, and for inducing arterial thrombosis after atherosclerotic plaque rupture. However, emerging evidence suggests a role for TF-coated microparticles in VTE. The exact mechanism and source of TF is still unclear [39]. Regardless of which cell specifically expresses TF in venous thrombosis, the lack of PAR4 showed a significantly increased clot formation time (CFT) in ROTEM with TF-mediated activation (EXTEM) (**Figure 4C**). In contrast, the clot formation time in blood from PAR4-P322L mice was identical to wild-type mice. This is likely due to the subtlety of the mutation and multitude of factors that contribute to clot formation in TF-mediated ROTEM in whole blood. The increased clot formation time in the PAR4^-/-^ mice suggests PAR4 promotes the early steps of clot formation but does not drive initiation of coagulation, overall clot stability, or clot firmness. This also reinforces the idea that reducing PAR4 activity therapeutically can effectively decrease venous thrombosis without significantly hindering primary hemostasis.The sharp decrease in the alpha angle also suggests a role for PAR4 in thrombin generation. Since procoagulant platelets are essential for mediating thrombin generation in hemostasis and thrombosis, impaired thrombin generation may be central to the mechanism for the decreased size and incidence of venous thrombi in our mutant mice.

Platelets promote coagulation by directly supporting thrombin activation and fibrin generation [7]. When platelets are exposed to a high concentration of agonist(s), as in vascular damage or thrombosis, they express phosphatidylserine on their surface [4,5]. This surface acts as a scaffold for the prothrombinase complex to assemble and generate thrombin [6,40]. Thrombin’s major function is to mediate fibrinogen conversion to fibrin, an insoluble protein that aggregates and stabilizes blood clots [19,41]. Thrombin also activates cells involved in thrombosis, including platelets, through protease activated receptors (PARs) [10]. When platelets are simultaneously activated *ex vivo* with a PAR4 agonist and a GPVI agonist, they develop a procoagulant phenotype expressing phosphatidylserine on their membrane [4]. Here, we showed that platelets from PAR4-322^L/L^ and PAR4^-/-^ mice had significantly less phosphatidylserine expression after simultaneous stimulation of PAR4 and GPVI (**Figure 5**).

Platelet-mediated thrombin generation also decreased as PAR4 reactivity decreased (**Figure 4A-C**). Altogether, we showed that as PAR4 reactivity decreases, platelet procoagulant activity also decreases providing a mechanism linking PAR4 to the thrombin-mediated initiation events of venous thrombosis such as activation coagulation cascade, immune cells, and endothelial cells.

In the early stages of venous thrombosis, platelets are recruited to the site where they associate with neutrophils and trigger TF expression, microparticle release, and NETosis [9,30]. This association is predominantly mediated by platelet P-selectin and neutrophil PSGL-1 [9,30]. Additionally, procoagulant platelets have characteristically high levels of P-selectin on their surface, and are known to preferentially bind to neutrophils [5,29,37]. Previously, we have shown that platelets from PAR4-322^P/L^ and PAR4-322^L/L^ mice require 2 and 8-fold higher concentration of PAR4-AP (AYPGKF), respectively, than wild-types to express their maximum level of P-selectin [24]. Here, platelet-neutrophil complexes were reduced concomitantly with decreased PAR4 reactivity (**Figure 7**). Complexes were unable to form from PAR4-322^L/L^ mice even at 400 μM, which triggers a full response from PAR4-322^P/P^ (wild-type). Taken with our previous data, PAR4-mediated platelet-neutrophil interactions are predominantly driven by PAR4 on platelets [24]. A direct role of PAR4 signaling on neutrophils cannot be ruled out, however, in our hands we have not observed PAR4-mediated signaling on neutrophils.

Here we show that the PAR4-P322L mutation negatively impacts venous thrombosis and platelet procoagulant activity. Our ex vivo experiments were consistent with previous observations that PAR4-322^L/L^ platelets have a more dramatic phenotype than heterozygous PAR4-322^P/L^ platelets. This gene-dosage effect implies that more leucines at PAR4-322 have a greater negative impact on platelet response to PAR4 agonists like thrombin. Conversely, our data from IVC stasis experiments show that the PAR4-322^P/L^ mice instead have a more dramatic phenotype than the PAR4-322^L/L^ mice, which are more similar to wild-types. Previously, experiments in arterial thrombosis showed a similar trend where PAR4-322^P/L^ mice, specifically females, had a more dramatic phenotype than their 322^L/L^ littermates [24]. An important distinction is that this switch was only seen in females, and here we only use males in our in vivo studies due to anatomical differences. It is also important to remember here that our mutation is global, and it impacts PAR4 on endothelial cells as well. We have only investigated the impact of the mutation of platelets specifically in this process, and it is likely that the PAR4-P322L mutation is also decreasing endothelial cell activation in response to thrombin.

PAR4 antagonists are currently not used therapeutically to prevent venous thrombosis. With our growing knowledge of how platelets promote venous thromboembolism, it is worth exploring PAR4 as a target. Venous clots are predominantly composed of red blood cells, fibrin, and neutrophils [32]. While platelets do not make up a large volume of the final thrombus, these small cells are vital to the beginning stages of thrombosis. Platelet participation in VTE has been known for over 50 years. In 1965, Hirsh and McBride reported that platelets from patients with recurrent VTE showed greater adhesive properties [42]. Platelets drive VTE by binding to damaged endothelium where they activate and promote thrombin generation and fibrin formation [30]. Here, we have shown that mice with the PAR4-P322L mutation have reduced venous thrombi size and incidence. This recapitulates clinical observations that humans with a leucine at PAR4-P310 are at a reduced risk of venous thromboembolism. We have also directly associated the PAR4-P322L mutation with reduced platelet procoagulant activity and complex formation with neutrophils, both of which promote thrombosis.Importantly, the complete lack of PAR4 delays clot formation but does not impair clot stability in ROTEM. Combined with our data that reduced PAR4 function negatively impacts both arterial and venous thrombosis, this highlights PAR4’s therapeutic potential to reduce thrombosis without impairing hemostasis.

## Author Contributions

Study Design: EK, ASG, MN Data Collection:EK, JC, NL, DD Data Analysis: EK, JC, NL, DD, MN Drafting Manuscript: EK, MN Critical Revisions: JG, NL, GH, DD, ASG

